# Autophagy regulates tumor growth and metastasis

**DOI:** 10.1101/2023.10.31.564991

**Authors:** Lei Qiang, Baozhong Zhao, Mei Ming, Ning Wang, Tong-Chuan He, Seungmin Hwang, Andrew Thorburn, Yu-Ying He

## Abstract

The role of autophagy in tumorigenesis and tumor metastasis remains poorly understood. Here we show that inhibition of autophagy stabilizes the transcription factor Twist1 through Sequestosome-1 (SQSTM1, also known as p62) and thus increases cell proliferation, migration, and epithelial-mesenchymal transition (EMT) in tumor development and metastasis. Inhibition of autophagy or p62 overexpression blocks Twist1 protein degradation in the proteasomes, while p62 inhibition enhances it. SQSTM1/p62 interacts with Twist1 via the UBA domain of p62, in a Twist1-ubiquitination-dependent manner. Lysine 175 in Twist1 is critical for Twist1 ubiquitination, degradation, and SQSTM1/p62 interaction. For squamous skin cancer and melanoma cells that express Twist1, SQSTM1/p62 increases tumor growth and metastasis in mice. Together, our results identified Twist1 as a key downstream protein for autophagy and suggest a critical role of the autophagy/p62/Twist1 axis in cancer development and metastasis.

## INTRODUCTION

Twist-related protein 1 (Twist1), also known as class A basic helix–loop–helix protein 38 (bHLHa38), is a basic helix-loop-helix (bHLH) transcription factor and is among the crucial factors regulating early embryonic development, carcinogenesis, and cancer metastasis (1–4). It suppresses E-cadherin-mediated cell-cell adhesion and promotes epithelial-mesenchymal transition (EMT) (3), and cell proliferation (4). Intriguingly, Twist1 is structurally unrelated to other EMT proteins such as Slug and Snail. Recent reports have shown that Twist1 is a short-lived protein degraded via the proteasomes (5). However, how Twist1 is regulated is not fully elucidated.

Autophagy (specifically macroautophagy) is a cellular self-eating system by which proteins, organelles, and other cellular materials are delivered for degradation in lysosomes (6, 7). Dysregulation of autophagy is observed in a number of diseases, including cardiovascular diseases, metabolic diseases, infection, neurodegeneration, and cancer (7–9). Sequestosome-1 (SQSTM1, also known as p62, hereafter ‘p62’) is an autophagy adaptor and substrate, linking ubiquitination with autophagy (10, 11). Previous reports have demonstrated the tumor-promoting role of p62 (12, 13) through regulating the critical transcription factors NF-kappaB (14, 15) and NRF2 (16–18). In addition, p62 expression is increased by Ras in tumorigenesis (15). Despite these progresses, the functions of autophagy and p62 in cancer pathogenesis remain poorly understood. In this study, we show that autophagy and p62 regulates EMT and Twist1 in tumor growth and metastasis.

## RESULTS

### Inhibition of autophagy reduces the expression of E-cadherin and increases cell migration, invasion, and proliferation

We first assessed the effect of autophagy inhibition on cellular malignant traits. Knockout (KO) of the essential autophagy genes Atg3, Atg5, Atg9 and Atg12 inhibited the conversion of LC3-I to LC3-II and caused p62 accumulation (Figure 1A), confirming autophagy inhibition. Autophagy inhibition reduced the expression of the negative regulator of cell proliferation and migration E-cadherin (Figure 1A), and increased cell invasion (Figure 1B), migration (Figure 1C), and cell proliferation (Figure 1D). In human skin cancer, well-differentiated (WD) and poorly-differentiated (PD) squamous cell carcinoma (SCC) showed increased p62 levels and decreased E-cadherin levels, as compared with normal human skin (Figure 1E). These findings indicated that inhibition of autophagy induces p62 accumulation, reduces E-cadherin level, and facilitates the oncogenic traits of cells and implicated that p62 may be a tumor-promoting factor associated with the decrease of E-cadherin expression in human SCCs.

**Figure 1.**
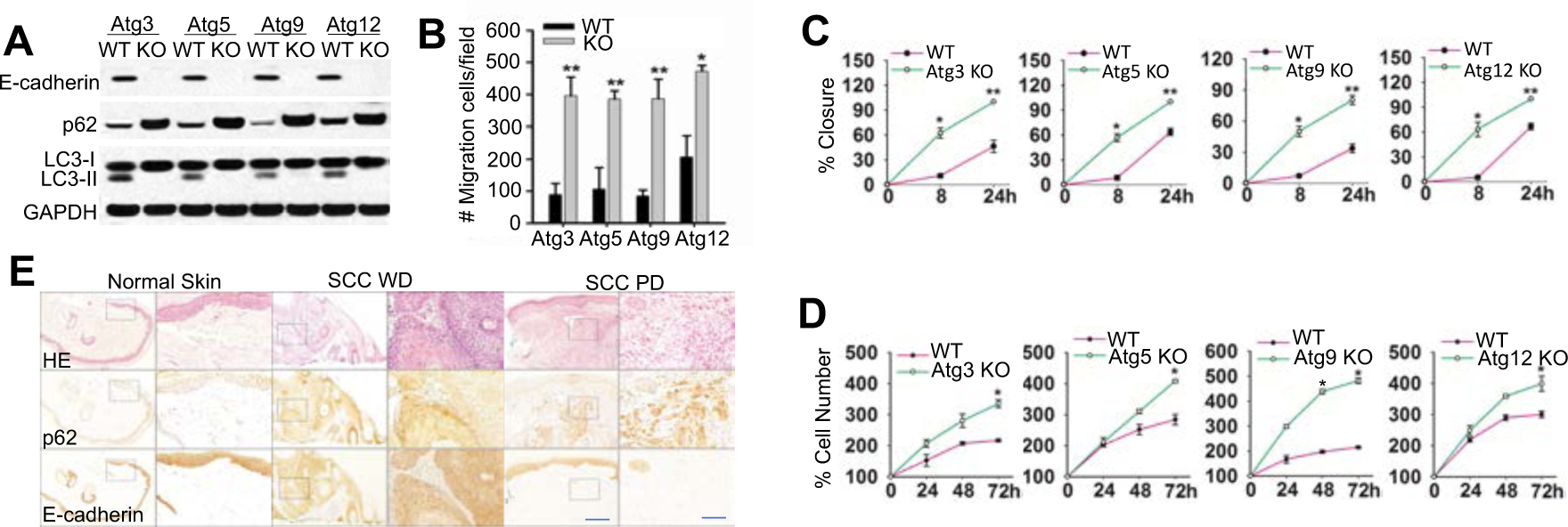
Effect of autophagy inhibition on cell migration, invasion, and proliferation. (A) Immunoblotting in cells with or without knockout (KO) of Atg3, Atg5, Atg9 or Atg12. (B) Transwell assay of cell invasion in cells as in A. (C) A radius migration assay of cells as in A. (D) Cell proliferation analysis of cells as in A. Data were obtained from three independent experiments (mean±S.D.). n=3. *, *P*<0.05; **, *P*<0.01, versus the WT group). (E) Representative images for histological and immunohistochemical analysis of p62 and E-cadherin (brown) in normal human skin (n=14), well-differentiated human squamous cell carcinoma (SCC WD) (n=25), and poorly differentiated human SCC (SCC PD) (n=10). Black squares present the region shown with higher magnification. Scale bar, 200 μm and 50 μm for left and right panels, respectively.

### Inhibition of autophagy suppresses the degradation of Twist1 protein

To assess how autophagy regulates E-cadherin expression, we analyzed whether inhibiting autophagy altered E-cadherin protein stability. E-cadherin levels was not affected by either CHX or MG132 in either WT or Atg5 KO cells (Figure S1A), suggesting that E-cadherin expression is regulated by autophagy at the mRNA level. Indeed Atg5 deletion reduced the E-cadherin mRNA level (Figure S1B), and the transcriptional activity of the wild-type (WT) E-cadherin promoter but not the E-Box mutant promoter (Figure 2A). These data demonstrated that inhibition of autophagy suppresses E-cadherin expression at the transcriptional level.

**Figure 2.**
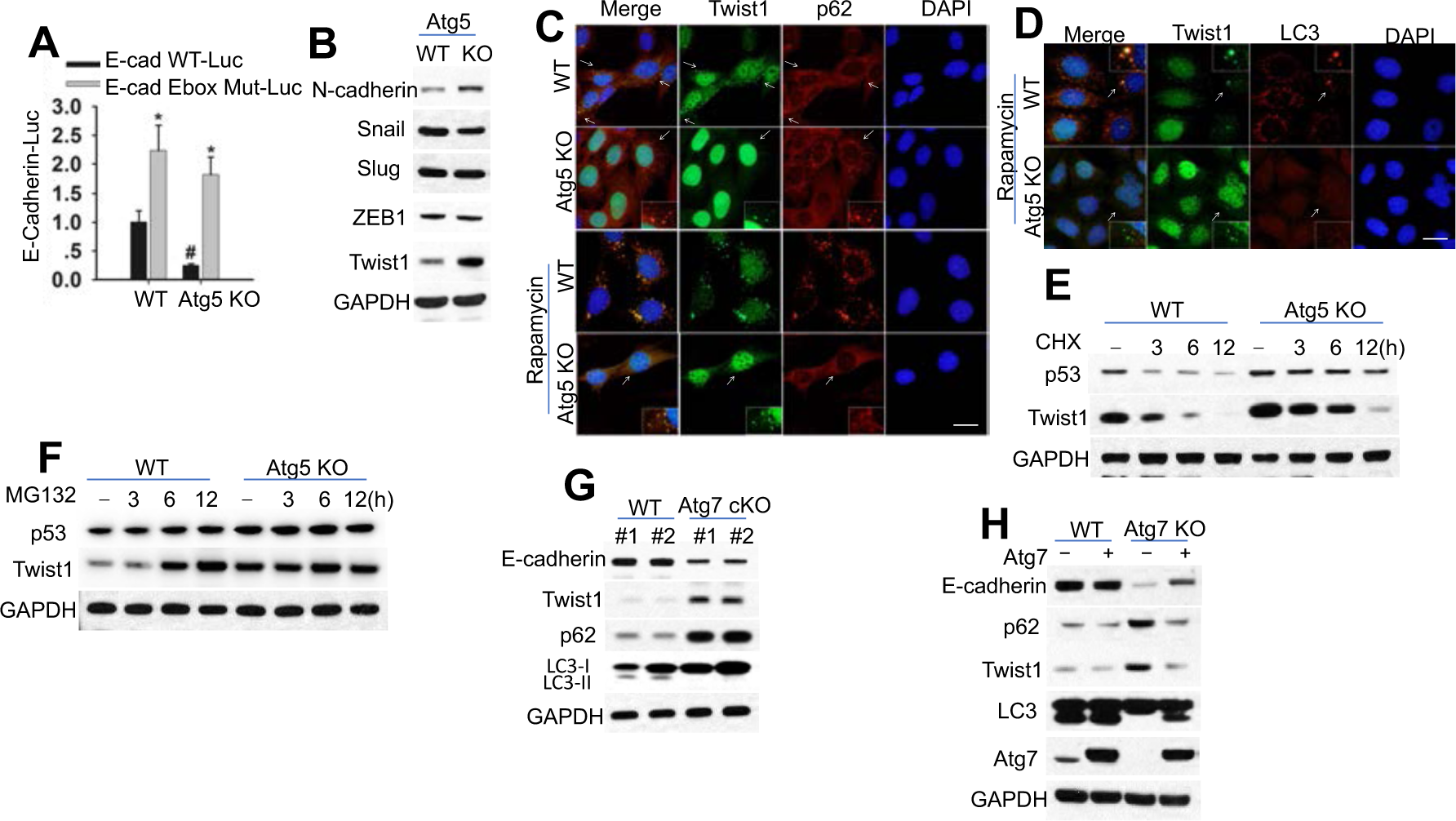
Autophagy inhibition suppresses Twist1 degradation. (A) Luciferase reporter assay of the E-cadherin promoter with an intact (E-cad (−108)-WT-Luc) or mutated (E-cad (−108) Ebox Mut-Luc) Ebox site in cells with or without Atg5 knockout. (B) Immunoblotting in cells with or without Atg5 knockout. (C-D) Immunofluorescence assay of p62/Twist1 (C) or LC3/Twist1 (D) in cells with or without Atg5 knockout treated with or without rapamycin (500 nM) for 6 h. Scale bar, 10 μm. DAPI is used as the nuclear counterstain. (E-F) Immunoblotting in cells with or without Atg5 knockout treated with or without CHX (100 μg/ml, E) or MG132 (10 μM, F) over a time course. (F) Immunoblotting in mouse skin with or without Atg7 knockout. (H) Immunoblotting in cells with or without Atg7 knockout in combination with Atg7 overexpression.

A number of suppressors for E-cadherin transcription have been identified, including Snail, Slug, ZEB-1 and Twist1, which interact with the proximal E-boxes of the E-cadherin promoter (19–21). Indeed, we found that Atg5 deletion increases the protein level of Twist1 while it has no effect on Snail, Slug, and ZEB1 (Figure 2B). Atg5 deletion increased of N-cadherin levels (Figure 2B) and the TCF/LEF activity (Figure S1C), a downstream signaling inhibited by E-cadherin (22). Autophagy inhibition increased Twist1, while p62 deletion decreased it (Figure S1D). However, autophagy deficiencies did not affect Twist1 mRNA level (Figure S1E). Twist1 overexpression suppressed E-cadherin expression (Figure S1F), Supporting the role of Twist1 in inhibiting E-cadherin expression. In cells with or without Atg5 deletion, p62 and Twist1 colocalized in punctate cellular structures independent of rapamycin treatment (Figure 2C). However, only WT cells but not in Atg5 KO cells showed colocalization of Twist1 with LC3 after rapamycin treatment (Figure 2D). It is worth noting that autophagy inhibition increased the protein stability of Twist1, but did not completely block Twist1 degradation (Figure 2E), suggesting that Twist1 is degraded through multiple systems, including autophagy and proteasomes. Indeed, MG132 increased Twist1 abundance in WT cells but not in autophagy-inhibited cells (Figure 2F, S1G). These results demonstrate that Twist1 is degraded via both autophagy and proteasomes.

To validate the regulation of Twist1 by autophagy in vivo in mice, we assesses the protein levels of E-cadherin, p62, and Twist1 in the mouse epidermis with wild-type (WT, K14Cre;Atg7+/+) and skin-specific deletion of Atg7 (K14Cre;Atg7flox/flox, Atg7 cKO). Epidermal Atg7 deletion increased the levels of both p62 and reduced the level of E-cadherin (Figure 2G, and S1H). Overexpression of Atg7 rescued the effect of Atg7 deletion (Figure 2H).

### p62 regulates Twist1 abundance independent of autophagy

To determine how p62 regulates Twist1, we analyzed the effect of p62 manipulation on Twist1 levels. p62 knockodwn reduced the Twist1 protein level in cells with or without autophagy inhibition (Figure 3A-B), while p62 overexpression up-regulate Twist1 (Figure 3C), supporting a role of p62 in regulating Twist1 independent of autophagy. p62 overexpression reversed the effect of p62 deletion (Figure 3D). p62 overexpression increased Twist1 protein stability (Figure 3E), while it did not affect the stability of Twist2 (Figure S2A), another member in the Twist subfamily of bHLH proteins (4). These results indicate that p62 increases Twist1 protein stability at least in part by inhibiting its degradation via the proteasomes.

**Figure 3.**
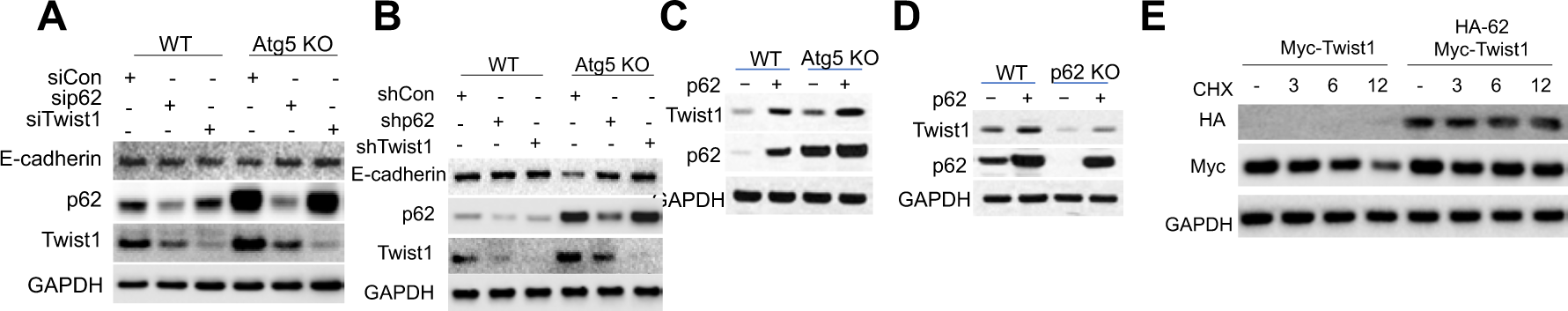
Regulation of Twist1 by p62. (A) Immunoblotting in cells with or without Atg5 knockout in combination with or without siRNA knockdown of p62 or Twist1. (B) Immunoblotting in cells with or without Atg5 knockout in combination with or without shRNA knockdown of p62 or Twist1. (C) Immunoblotting in cells with or without Atg5 knockout in combination with or without p62 overexpression. (D) Immunoblotting in cells with or without p62 knockout in combination with or without p62 overexpression. (E) Immunoblotting in 293T cells with or without overexpression of Myc-Twist1 and HA-p62 treated with or without CHX (100 μg/ml) over a time course.

### p62 interacts with Twist1

Next we assessed whether p62 interacts with Twist1. Indeed we found that endogenous Twist1 interacted with p62 (Figure 4A). Atg5 deletion increased the total and Twist1-interacting p62 level, while it decreased the Twist1-interacting level of Rad23B (Figure 4A), a protein delivering substrates to proteasomes (23–25). Rad23B overexpression decreased the Twist1 protein levels in cells with or without Atg5 deletion (Figure 4B), implicating a critical role of competitive binding of Twist1 to either RAD23B or p62 in regulating Twist1 protein stability.

**Figure 4.**
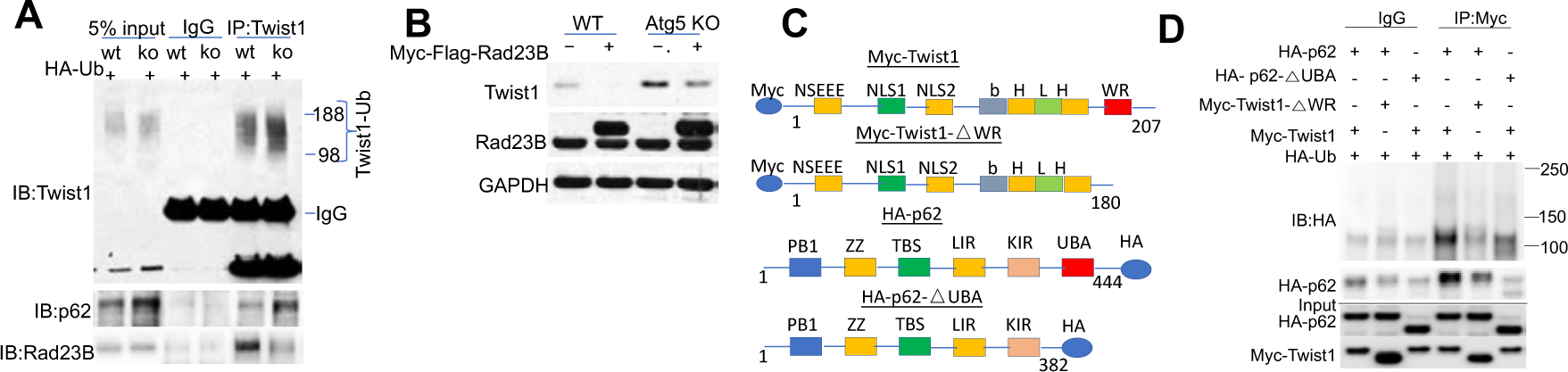
p62 interacts with Twist1. (A) Co-immunoprecipitation (Co-IP) analysis of Twist1 interaction with p62 or Rad23B in cells with or without Atg5 deletion. (B) Immunoblotting in cells with or without Atg5 deletion in combination with overexpression of Myc-Flag-Rad23B or empty vector (EV). (C) Schematic for the constructs with or without individual domain deletions for p62 and Twist1. (D) Co-IP analysis of the interaction between p62 or p62-△UBA and Twist1 or Twist1-△WR in 293T cells. Molecular weight in kDa is marked (A and D).

To determine how Twist1 degradation is regulated, we assessed the role of the WR domain of Twist1, which is essential for ubiquitination and degradation of Twist1 in xenopus (5). We generated a Twist1 mutant with WR deletion (Myc-Twist1-△WR) (Figure 4C). Indeed, WR deletion stabilized Twist1 (Figure S2B-D). In addition, we found that the WR domain was required for Twist1 ubiquitination and Twist1-p62 interaction (Figure 4D). p62 has been reported to interact with ubiquitinated protein targets via its UBA domain (26). Indeed the UBA domain of p62 was required for the interaction of p62 with ubiquitinated Twist1 (Figure 4D). These data indicate that p62 interacts ubiquitinated Twist1, which requires the WR domain in Twist1 and the UBA domain in p62.

### Role of lysine 175 in p62-regulated Twist1 protein stability

Next we set out to identify the critical lysine sites for Twist1 ubiquitination. To do so, we created several lysine mutants (lysine to arginine, K->R) (Figure S3A). It is worthwhile to note that all the lysine sites are outside of Twist1 WR domain (Figure S3A). Among all the mutants, The K175R mutant demonstrated higher protein levels than WT Twist1 and other mutants, mimicking the WR deletion mutant (Figure 5A). K175R mutation also enhanced Twist1 protein stability (Figure 5B), demonstrating that K175 is critical for the degradation of Twist1. In addition, we found that K175 is also critical for Twist1 ubiquitination (Figure 5C). K175R mutation inhibited Twist1 interaction with p62 and colocalization with p62, mimicking WR deletion (Figure 5C-D). K175R mutant showed increased protein Twist1 abundance and increased E-cadherin inhibition (Figure 5E). p62 only affected wild-type Twist1, but not the K175R mutant (Figure 5E). In contrast, despite upregulated protein level, the WR deletion mutant did not affect the E-cadherin level (Figure 5E), supporting the importance of the WR domain in Twist1 function (27). In addition to K175, we also determined the importance of other lysines in Twist1. Mutations of these sites demonstrated that both the K73-76-77 sites and the K137 site are also critical for Twist1 degradation (Figure S3B). However, Twist1 stability was not affected by mutation of the individual sites alone (Figure S3C). These findings demonstrate that K175 is required for ubiquitination, degradation, and interaction of Twist1 with p62 (Figure 5F).

**Figure 5.**
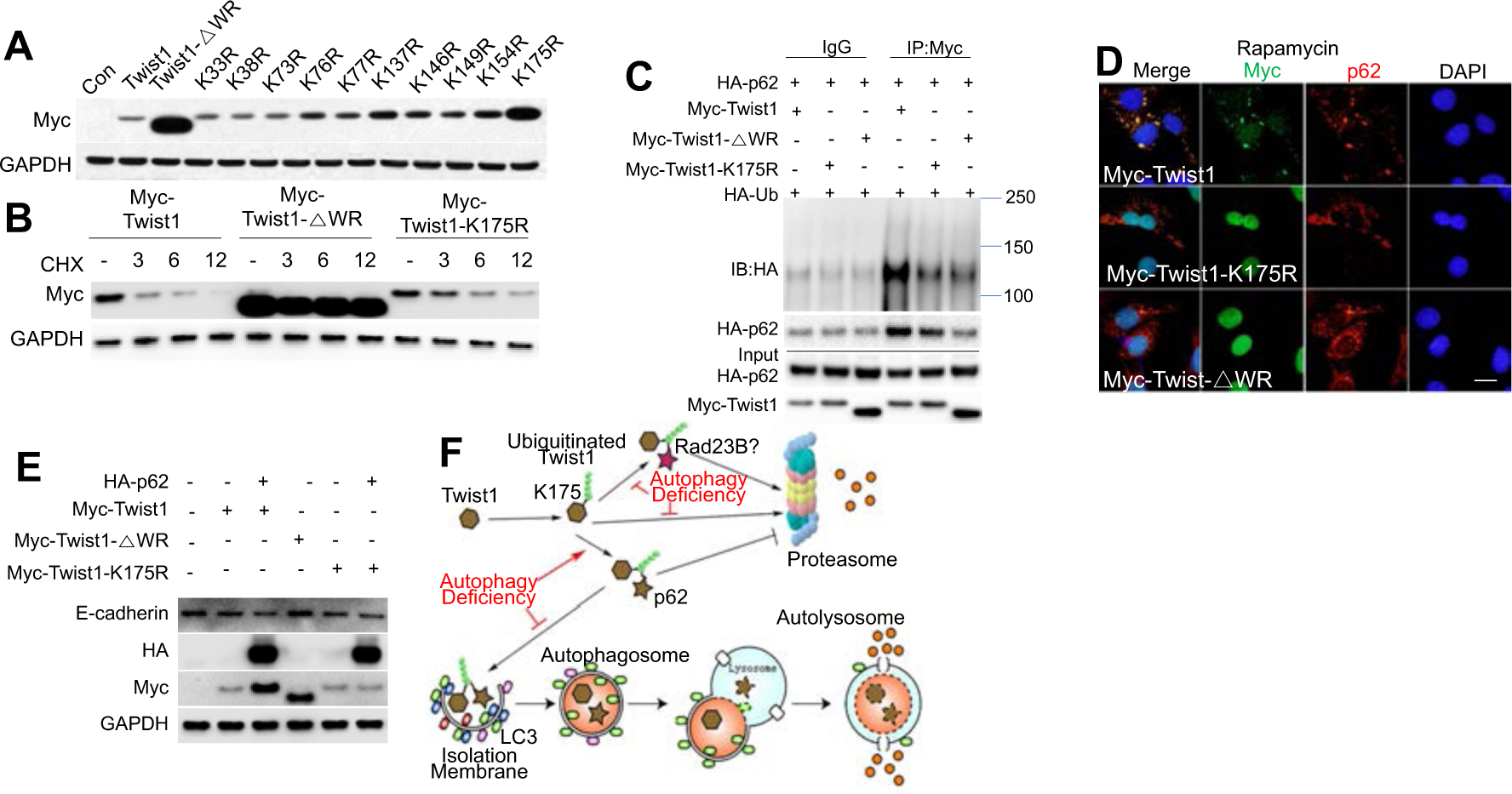
Role of K175 in ubiquitination, degradation, and interaction of Twist1 with p62. (A) Immunoblotting in 293T cells overexpressed with vector (Con), WT, or Twist1 mutants. (B) Immunoblotting in 293T cells transfected with WT or Twist1 mutants followed by treatment with or without with CHX (100 μg/ml) over a time course. (C) Co-IP analysis of ubiquitination and interaction of WT and mutant Twist1 with p62. (D) Immunofluorescence assay of p62 and Myc (Twist1) in WT MEF cells transfected with WT or mutant Twist1 following treatment with rapamycin (500 nM) for 6 h. Scale bar, 10 μm. DAPI is used as the nuclear counterstain. (E) Immunoblotting in WT MEF cells with or without overexpression of WT or mutant Twist1 in combination with p62 overexpression. (F) Schematic summary for Twist1 stabilization by p62.

### The role of the p62/Twist1 axis tumor growth and metastasis in mice

To address the importance of the regulation of Twist1 b p62 in cancer, we analyzed the role of p62 in vitro and in vivo. In Twist1-absent A431 cancer cells, overexpression of p62 did not affect on E-cadherin expression, cell proliferation, or migration (Figure 6A-D). Upon forced Twist1 overexpression, p62 overexpression up-regulated Twist1, decreased E-cadherin and the E-cadherin-β-catenin colocalization at the plasma membrane, which mediates E-cadherin-loss-regulated migration (28), and enhanced cell proliferation and migration (Figure 6A-D, S4A). In mice, when Twist1 was absent, p62 overexpression di not affect either tumor growth (Figure 6E) or metastasis (no metastasis detected for either control or p62-added A431 cells). However, when Twist1 was added, p62 overexpression enhanced both tumor growth and metastasis (Figure 6E-F). p62/Twist1 overexpression decreased E-cadherin expression, increased the number of Ki67-positive cells, and resulted in a loss of tumor boundary (Figure S4B). In Twist1-expresing human A375 melanoma cells, p62 overexpression enhanced both the Twist1 level (Figure 6G) and tumor growth in nude mice (Figure 6H), while p62 inhibition decreased them (Figure 6I-J). Atg7 inhibition resulted in an increase in the levels of both Twist1 and p62 and tumor growth in nude mice (Figure 6I-J). These findings demonstrate that Twist1 regulation by p62 promotes tumor growth and metastasis in vivo.

**Figure 6.**
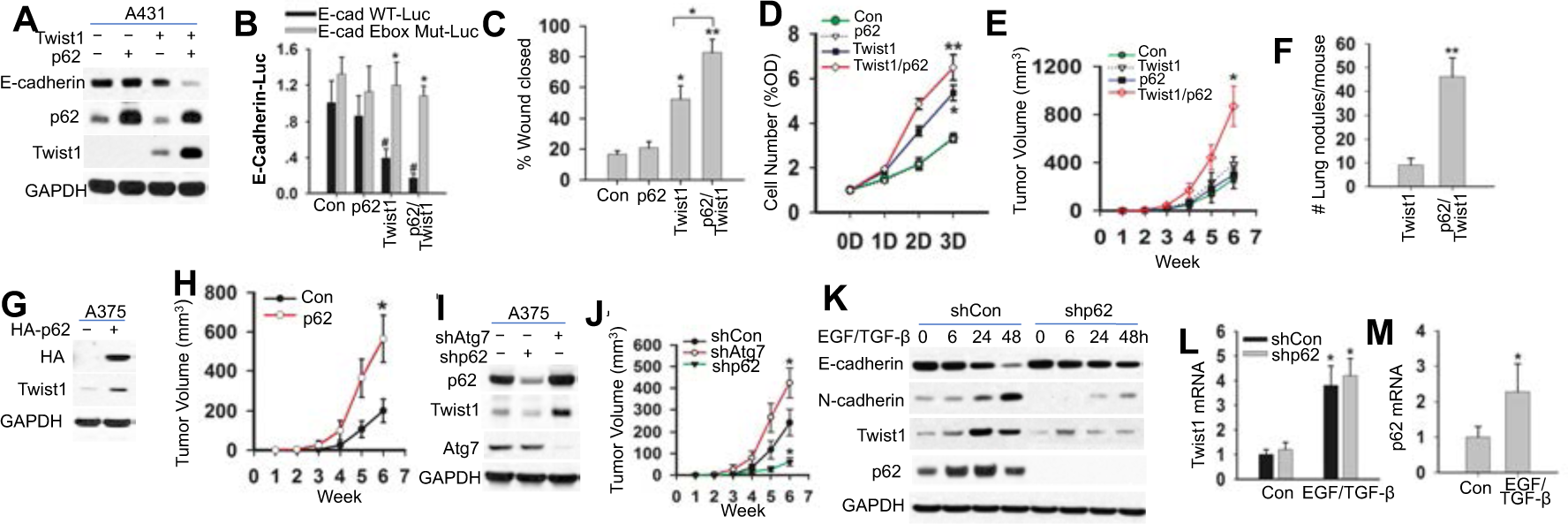
Role of the p62/Twist1 axis in tumor growth and metastasis. (A) Immunoblotting in A431 cells with or without overexpression of p62 and/or Twist1. (B) Luciferase reporter assay of the E-cadherin promoter with an intact (E-cad (−108) WT-Luc) or mutated (E-cad (−108) Mut-Luc) Ebox site in cells as in A. (C) Wound healing assay of cell migration in cells as in A. (D) MTS assay of cells proliferation of cells as in A. (E) Tumor growth from cells as in A. (F) Number of lung tumor nodules per at 14 weeks following injection. (G) Immunoblotting in A375 cells with or without p62 overexpression. (H) Tumor growth from cells as in G. (I) Immunoblotting in A375 cells transfected with or without knockdown of Atg7 or p62. (J) Tumor growth from cells as in I. (K) Immunoblotting in HaCaT cells with or without p62 knockdown and treated with or without EGF/TGF-β)over a time course. (L) qRT-PCR analysis of Twist1 in HaCaT as in K treated with EGF/TGF-β for 48 h. (M) qRT-PCR analysis of p62. Data are shown from three independent experiments (mean±S.D.). n=3. *, *P*<0.05; compared with WT Con cells (B); #, *P*<0.05; compared with the WT group (B); *, *P*<0.05; compared with the Con, A431-p62, and A431-Twist1 groups (E); **, *P*<0.01; compared with the A431-Twist1 group (F).

To determine whther the p62/Twist1 axis regulates EMT, we treated the low-Twist1-expressing HaCaT cells with EGF and TGF-β for 48h as described previously (29). In control (Con) HaCaT cells, EGF/TGF-β induced EMT, inhibited the expression of E-cadherin, and promoted the expression of N-cadherin, p62, and Twist1 (Figure 6K-M, S4C). EGF/TGF-β-induced p62 expression preceded the Twist1 induction (Figure 6K). p62 knockdown reduced the up-regulation of Twist1 and suppression of E-cadherin, and suppressed EGF/TGF-β-induced EMT (Figure 6K, S4C). However, p62 knockdown did not afffect the mRNA level of Twist1 (Figure 6L). These findings demonstrate that p62 is critical for Twist1 accumulation in EMT.

## DISCUSSION

The mechanism for the regulation of cancer development and metastasis remains poorly understood. Here we show that the autophagy adaptor protein p62 interacts with Twist1 and thus promote Twist1 protein stabilization to promote tumor growth and metastasis. Twist1 seems to be degraded via both the autophagy and proteasomal pathways (Figure 5F). p62 competes with Rad23B in binding with Twist1, which may inhibit Twist1 delivery by Rad23B to proteasome for degradation (23–25) (Figure 5F). As a result, p62 enhances Twist1-mediate EMT. These findings provided a new mechanism by which Twist1 is regulated in tumor promotion and progression.

In healthy adult tissues, Twist1 expression was largely undetectable; however it is up-regulated in a number of cancers (30–34). Recently autophagy induction by death effector domain-containing DNA-binding protein (DEDD) increases the degradation both Twist and Snail through autophagy (35). Furthermore, p62 interacts with ubiquitinated proteins to deliver them to proteasomes for degradation (26), including the protein Tau (36). In contrast, our results indicate that p62 accumulation by autophagy inhibition promote the accumulation of Twist1, but not other EMT factors, and that p62 suppresses the degradation of Twist1 via both the autophagy and proteasome pathways. We demonstrated that p62 interacts with Twist1 to inhibit Twist1 interaction with Rad23B, likely leading to reduced Twist1 delivery to the proteasomes. These different mechanisms can be resulted from the cell-type-dependent effect of Twist1 degradaion under different contexts. Indeed p62 has been shown to interact with ubiquitinated p53, thus blocking its degradation via the proteasomes at least in part in part by inhibiting p53 interaction with the ubiquitin-dependent chaperone protein p97/VCP (37). Future investigations will characterize the specific role of p62 in modulating degradation of different substrates and characterize the additional role f p62 on Twist1 fucntion, such as regulating Twist1 activity.

In addition to autophagy deficiency, we found that p62 also promotes Twist1 accumulation in EMT. It is possible that the p62/Twist1 axis may also have an active role in Ras-driven cancers such as lung and pancreatic cancers, as Ras induces p62 expression (15). Future investigation is warranted to elucidate the importance of the p62/Twist1 interaction in cancer progression as well in cancer initiation. Besides Twist1, recent reports also demonstrated that autophagy also controls RhoA signaling through p62 (38, 39). Future investigations are needed to assess the importance of the p62/Twist1 axis in clinically relevant genetic mouse tumor models.

## METHODS

### Human Samples

All human specimens were analyzed after the approval by the University of Chicago Institutional Review Board. Skin samples were obtained from the archives in the tissue bank from the Section of Dermatology, Department of Medicine, University of Chicago. Non–sun-exposed normal skin samples and squamous skin carcinomas were analyzed for histology and for E-cadherin and p62 protein levels by immunohistochemical staining.

### Cell culture

WT, Atg5 KO MEF cells (obtained from Dr. Mizushima), Atg3 KO, Atg7 KO, Atg9 KO, Atg12 KO, p62 KO, Control and Atg14 cKO MEF cells, doxycycline-inducible Atg12 knockdown (KD) 4T1 cells, HaCaT cells (kindly provided by Dr. Fusenig), HEK293T cells, A431 cells, and A375 cells were maintained in a monolayer culture in 95% air/5% CO_2_ at 37℃ in Dulbecco’s modified Eagle’s medium (DMEM) supplemented with 10% fetal bovine serum (FBS), 100 units per mL penicillin, 100 μg per mL streptomycin (Invitrogen, Carlsbad, California), and 1% NEAA (Nonessential Amino Acids, Invitrogen, Carlsbad, California). For the EMT analysis, cells were first washed with PBS twice and then incubated with DMEM medium with or without EGF (10 ng/μl, R&D) and TGF-β (10 ng/μl, R&D) for 48 h.

### siRNA and plasmids transfection

MEF cells were transfected with negative control (NC) or siRNA (ON-TARGETplus SMARTpool, Dharmacon) targeting Twist1 or p62, using DharmaFECT 4 transfection reagent (Dharmacon, Lafayette, CO) according to the manufacturer’s instructions as described previously (40). HEK-293T and MEF cells (except for p62 KO MEF cells) were transfected with plasmids using X-tremeGENE 9 (Roche) as described as previously (41–43). A431 and A375 cells were transfected using Amaxa Nucleofector according to the manufacturer’s instructions as described previously (41, 42). Lentiviral vectors were used for overexpression of Atg7 and p62 and gene knockdown by shRNA. HEK293T cells were used as packaging cells for the lentiviral system. pLKO.1 vectors and packaging mix (psPAX2 and pMG2) were transfected into HEK293T cells. Two days after transfection, supernatants were used for cell infection. Polybrene was at 8 μg/ml for generating stable cell lines.

### Plasmids

Myc-Twist1 pcDNA3.1 and Flag-Twist1 pcDNA3.1 were kindly provided by Dr. Tony Firuli. Myc-Twist1 (HindIII/XbaI) from pcDNA3.1 vector was cloned to pMPB3 vector (modified piggyBac vector). HA-p62 pcDNA4 was kindly provided by Dr. Qing Zhong (Addgene plasmid 28027) (44). HA-Ubiquitin pcDNA3 was kindly provided by Dr. Edward T. H. Yeh (Addgene plasmid 18712)(45). PGL2 E-cad (108) WT-Luc and PGL2 E-cad (108) Ebox Mut-Luc were kindly provided by Dr. Eric R. Fearon (Addgene plasmids 19291 and 19290)(46). The TCF/lymphoid enhancer factor luciferase reporter (TOP) and its negative control (FOP) plasmids were kindly provided by Dr. Tong-Chuan He. Myc-Flag-Rad23b-pCMV, Flag-Twist2 and pCMV6 vector were purchased from OriGene Technologies. pLKO.1 shp62 (human) was purchased from Sigma. pLKO.1 shp62 (mouse) and pLKO.1 shTwist1 (mouse) were purchased from Dharmacon. pENTER vector, pHAGE con and pHAGE Atg7 (mouse) were kindly provided by Dr. Seungmin Hwang. pLenti CMV Dest vector was purchased from addgene. pLKO.1 shAtg7 (human) was kindly provided by Dr. Kimmelman.

### DNA constructs

Myc-Twist1-WR, the WR deletion mutant of Twist1, were created with the following primers: Forward-5’-<u>TACGACTCACTATAGGGAGACCC-3’ and</u> Reverse-5’-GTTCTAGACTAGCAGCTTGCCATCTTGGAGTCC-3’ from Myc-Twist1 pcDNA3.1.It was then subcloned (HindIII/XbaI) into pMPB3 vector. Domain deletion mutants of p62 (A, B, C, D, E and F) were generated using the primers C-Terminal sense 5’-GGAATTCTATGGTGCACCCCAATGTGATCTG-3’ and N-Terminal primer A 5’-GGAATTCTATGGTGCACCCCAATGTGATCTG-3’, primer B 5’-GGAATTC-TATGGGTCCACCAGGAAACTGGA-3’ and primer C 5’-GGAATTCTATGGAGTCGGA-TAACTGTTCAGG-3’; N-Terminal sense 5’-GGAATTCTATGGCGTCGCTCACCGTGAA-3’ and C-Terminal primer D 5’-GAGTGCGGCCGCACGGGTCCACTTCTTTTGAAG-3’, primer E 5’-GAGTGCGGCCGCAGCTTGGCCCTTCGGATTCT-3’ and primer F 5’-GAG-TGCGGCCGCGGCTTCTTTTCCCTCCGTGCT-3’ and subcloned (EcoR1/Not1) back into the HA-pcDNA4 vector. HA-p62 WT was generated using the primers N-Terminal sense 5’-GTGGTACCGCTATGGCGTCGCTCACCGTGAA-3’ and C-Terminal sense 5’-GAGGCGCGCCCTAAGCGTAATCTGGAACATCGT-3’ from HA-p62 pcDNA4 and subcloned (Kpn1/Asc1) into pMPB3 vector and pENTER vector. Generation of stable transfected cell lines using the piggyBac transposon system (pMPB3) were performed as described previously (47). pENTER p62 was recombined into pLenti CMV Dest vector using Gateway LR Clonase Enzyme Mix Kit according to the manufacturer’s instructions. All plasmids were confirmed by sequencing.

### Site directed mutagenesis

Mutations at lysine 33, 38, 73, 76, 77, 137, 146, 149, 154 and 175 of Twist1 in the pMPB3 vector were performed using the QuikChange XL site-directed mutagenesis kit accorinding to the manufacturer’s instructions (Stratagene), using the following primers. K33R 5’-CCGGCGAGCGGCAGGCGCGGG-3’ and K33R antisense 5’-CCCGCGCCTGCCGCTCGCCGG-3’; K38R 5’-GCGGGGCTCGCAGGAGAC-GCAGCAG-3’ and K38R antisense 5’-CTGCTGCGTCTCCTGCGAGCCCCGC-3’; K73R 5’-CCGGCCCAGGGCAGGCGCGGC-3’ and K73R antisense 5’-GCCGCGCCTGCCCTGGGCCGG-3’; K76R 5’-GGCAAGCGCGGCAGGAAATCTGCGGGC-3’ and K76R antisense 5’-GCCCGCAGAT-TTCCTGCCGCGCTTGCC-3’; K77R 5’-AGCGCGGCAAGAGATCTGCGGGCGG-3’ and K77R antisense 5’-CCGCCCGCAGATCTCTTGCCGCGCT-3’; K137R 5’-CCGCCCTGCGCAGGATCATCCCCAC-3’ and K137R antisense 5’-GTGGGGATGATCCTGCGCAGGGCGG-3’; K146R 5’-CACGCTGCCCTCG-GACAGGCTGAGCAAG-3’ and K146R antisense 5’-CTT-GCTCAGCCTGTCCGAGGGCAGCGTG-3’; K149R 5’-CTCGGACAAGCTGAGCAGGATTCAGACCCTCAAAC-3’ and K149R antisense 5’-GTTTGAGGGTCTGAATCCTGCTCAGCTTGTCCGAG-3’; K154R 5’-AGCAAGAT-TCAGACCCTCAGACTGGCGGCC-3’ and K154R antisense 5’-GGCCGCCAG-TCTGAGGGTCTGAATCTTGCT-3’; K175R 5’-CGAGCTGGACTCCAGGATGGCAA-GCTGCA-3’ and k175R antisense 5’-TGCAGCTTGCCATCCTGGAGTCCAGCTCG-3’; all mutants were confirmed by sequencing.

### Immunofluorescence

Immunofluorescence staining was performed as described previously (40, 48). Briefly, cells cultured on the coverslips were washed three times with PBS and fixed with 4% paraformaldehyde for 25 min. Cells were then permeabilized with cold 0.5% Triton-X-100 in PBS for 20 min at 4°C and washed with cold PBS. Cells were then blocked with 5% goat serum (Invitrogen, Carlsbad, California) (30 min, 37°C). Next cells were incubated at 4°C overnight with anti-E-cadherin (1:200; Cell Signaling Biotechnology), β-Catenin (1:200 BD Biosciences, Franklin Lakes, NJ), HA (1:200 Cell Signaling Biotechnology), Myc (1:200 Cell Signaling Biotechnology), LC3 (1:200 Cell Signaling Biotechnology), Twist1 (1:200 Abcam). or p62 (1:200 Progene Biotechnik, Heidelberg, Germany) antibody. After washing with PBS, cells were next incubated at 37°C for 1 h with Alexa Fluor 488 F (ab’) 2 fragments of goat anti-mouse IgG antibodies and Alexa Fluor 568 of goat anti-rabbit IgG antibodies. Cells were then fixed in Prolong Gold Antifade with DAPI (Invitrogen, Carlsbad, California) as a nuclear counterstain, followed by imaging under a fluorescence microscope (OlympusIX71, Japan).

### Immunoblotting

Immunoblotting was performed as described previously (49). The following antibodies were used: p53, Rad23b, GAPDH and HA-Tag (mouse) (Santa Cruz), E-cadherin, LC3I/II, HA-Tag (Rabbit), Myc-Tag (mouse), Myc-Tag (Rabbit), Snail, Slug, Atg7, Flag-Tag, N-cadherin, ZEB1 (Cell Signaling Technology), p62 (Progene Biotechnik, Heidelberg, Germany), and Twist1 (Abcam). Protein levels were measured using ImageJ and normalized against the protein level at t=0. Log_10_ of the percentage of the initial protein level was plotted versus time, and half-life (t_1⁄2_) was calculated from the log_10_ of 50%.

### Luciferase Reporter Assays

Cells were transfected with PGL2 E-cad (108) WT-Luc, PGL2 E-cad (108) Ebox Mut-Luc (kindly provided Dr. Eric R. Fearon, Addgene plasmids 19291 and 19290), and 0.025 μg of pRL-TK (Promega, used as a transfection efficiency control) using X-treme 9 Transfection Reagent (Roche) or Amaxa Nucleofector following the manufacturer’s protocols. Luciferase activity was analyzed as described previously (41).

### Immunohistochemical staining

Immunohistochemical staining of E-cadherin and Ki67 was performed by the Immunohistochemistry core facility at the University of Chicago. E-cadherin and Ki67 (Cell signaling) antibodies were used, following by visualization with the diaminobenzidine (DAB) method (brown color). To determine the contribution of brown signal due to melanin, hematoxylin and eosin (H&E) staining was also performed.

### Real-time qPCR

qPCR analysis were carried out using ABI7300 (Applied Biosystems, Foster City, CA) with the SYBR® Green PCR Master Mix (Applied Biosystems) as in our previous studies (50). The following primers were used: 5’-CAGGTCTCCTCATGGCTTTGC-3’ (forward), 5’-CTTCCGAAAAGAAGGCTGTCC-3’ (reverse) for the mouse E-cadherin gene; 5’-AG-GTCGGTGTGAACGGATTTG-3’ (forward) and 5’-TGTAGACCATGTAGTTGAGGTCA-3’ (reverse) for GAPDH; 5’-CAGAGAAGCCCATGGACAG −3’ (forward) and 5’-AGCTGCCTTGTACCCACATC −3’ (reverse) for the human p62 gene; 5’-CGGGAGTCCGCAGTCTTA −3’ (forward) and 5’-GCTTGAGGGTCTGAATCTTG −3’ (reverse) for the human Twist1 gene; and 5’-GCAGGACGTGTCCAGCTC-3’ (forward) and 5’-CTGGCTCTTCCTCGCTGTT-3’ (reverse) for the mouse Twist1 gene.

### Mouse samples and tumor growth and metastasis in nude mice

All animal procedures were approved by the University of Chicago Institutional Animal Care and Use Committee. K14-Cre mice were obtained from Jackson Laboratories and Atg7flox/flox mice were kindly provided by Masaaki Komatsu (Tokyo Metropolitan Institute of Medical Science, Tokyo, Japan).

Tumor cells (1×10^6^) were injected subcutaneously into the right flank of 6-week-old female nude mice (Harlan Sprague Dawley). Tumors were measured weekly. Tumor volume (TV) was analyzed as follows: TV (mm^3^) = d^2^×D/2 where d and D are the shortest and the longest diameters, respectively. For metastasis experiments, cells were tail vein injected as described previously (51). Lungs and other tissues with metastasis were collected for various analyses.

### Migration and invasion assay

For migration analysis, cells were plated onto Radius™ 24-Well Cell Migration Assay plates following the manufacturer’s protocol (Cell Biolabs, Inc. San Diego, CA). To analyze the invasion activity, the upper surface of the Transwell filters (8 μm pore size Millipore MultiScreen-MIC plates) was coated with Matrigel as described previously (52). Cells were cultured to be confluent and quiescent. Next the Gel Spot was removed within 5 min, and then the cells were populated the circular void space. Digital images of the gap closure were obtained with an inverted microscope (OlympusIX71, Japan). Images were assessed with the CellProfiler™ Cell Image Analysis Software.

### *In vitro* cell proliferation assay

Cell proliferation was measured using the MTS assay (Promega, Madison, WI, USA) following the manufacturer’s instructions.

### Statistical analyses

Statistical analyses were carried out using Prism 5 (GraphPad software, San Diego, CA, USA). Data were analyzed by Student’s t-test. A *P*-value of <0.05 was considered statistically significant.

## ACKNOWLEDGMENTS

We are grateful to Drs. Deborah Lang and Kay Macleod for their discussion and suggestions, and Dr. Ann Motten for a critical reading of the manuscript. We are grateful to all the investigators for kindly providing plasmids and mice as specifically indicated in the Methods section, and Terri Li at the HTRC core facility for immunohistochemistry. This work was supported by the NIH/NIEHS grant ES016936 (YYH), the American Cancer Society (ACS) grant RSG-13-078-01 (YYH), the University of Chicago Cancer Research Center (P30 CA014599), the CTSA (NIH UL1RR024999), and the University of Chicago Friends of Dermatology Endowment Fund.

---

**Figure S1, related to Figure 2.**
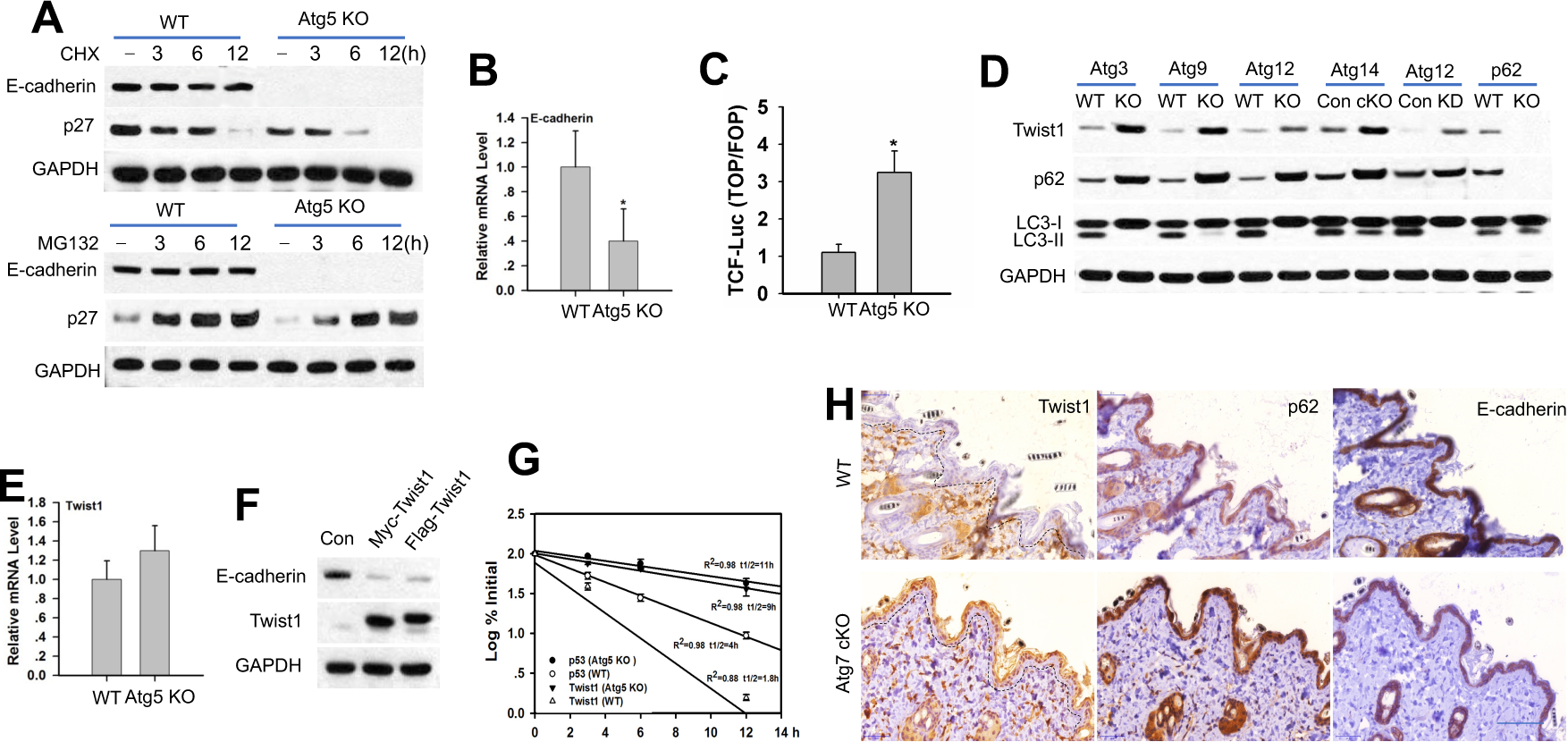
Autophagy regulates Twist1 protein degradation. (A) Immunoblotting in MEF cells with or without Atg5 deletion treated with cycloheximide (CHX 100 μg/ml) or MG132 (10 μM) over a time course as in Figure 2E-F. (B) qPCR analysis of the mRNA level of E-cadherin in MEF cells with or without Atg5 deletion. (C) Luciferase analysis of the TCF/lymphoid enhancer factor luciferase reporter (TOP) and its negative control (FOP) in MEF cells with or without Atg5 deletion. Data were shown from three independent experiments (mean±S.D.). n=3. *, p<0.05; **, p<0.01, compared with the Con group). (D) Immunoblotting in cells with or without deletion of autophagy genes. (E) qPCR analysis of Twist1 mRNA levels in MEF cells with or without Atg5 deletion. (F) Immunoblotting in MEF cells with or without Twist1 overexpression. (G) Quantification of the Twist1 protein levels in Figure 2E. Data were shown from three independent experiments (mean±S.D.). n=3. *, *P*<0.05 compared with the WT cells; #, *P*<0.05 compared with the WT promoter).

**Figure S2, related to Figure 4.**
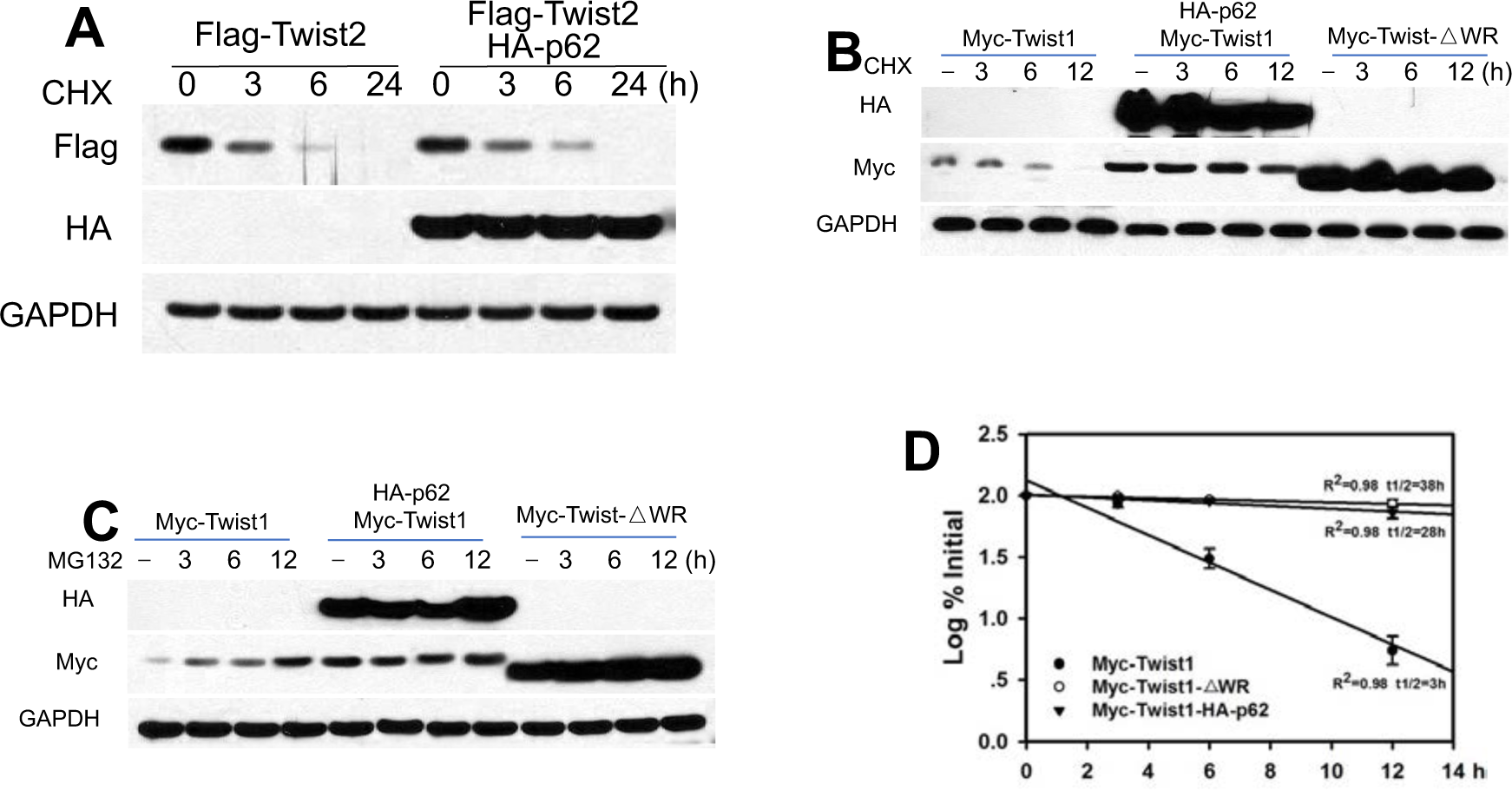
p62 interacts with Twist1. (A) Immunoblotting in 293T cells with or without overexpression of Twist2 and/or p62 treated with or without CHX (100 μg/ml) over a time course. (B-C) Immunoblotting in 293T cells expressing Myc-Twist1, Myc-Twist1/HA-p62, or Myc-Twist1-△WR and treated with CHX (100 μg/ml) (B) or MG132 (10 μM) (C) over a time course. (D) Quantification of B.

**Figure S3, related to Figure 5.**
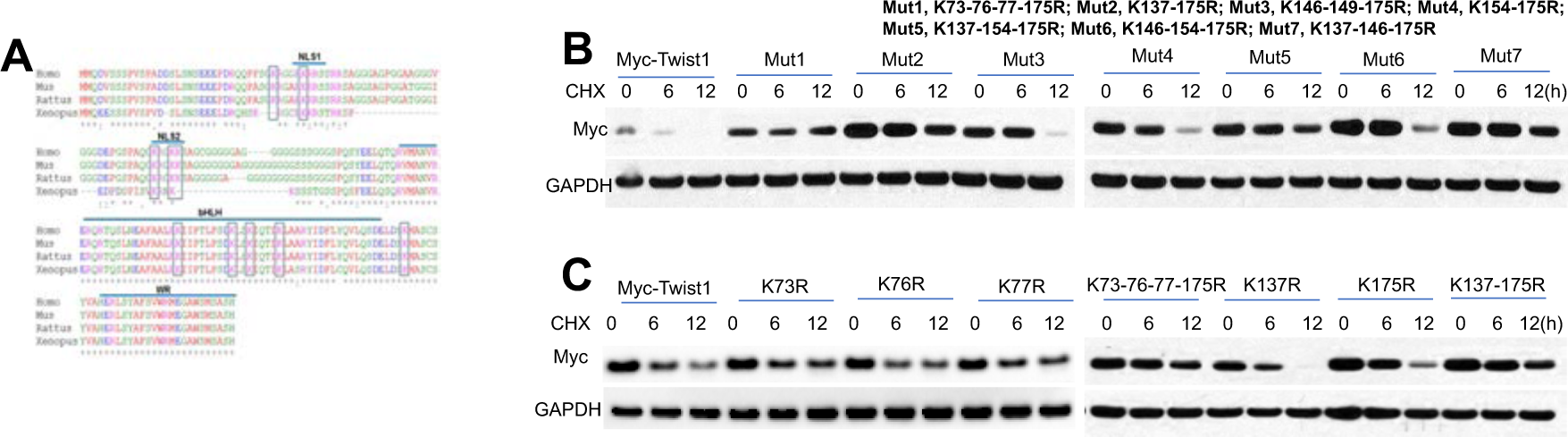
Lysine 175 of Twist1 is critical for ubiquitination, degradation, and interaction of Twist1 with p62. (A) Amino acid sequences across Human (Homo), Mouse (Mus), Rat (Rattus) and Xenopus. Lysine sites are indicated with rectangles. “*” indicates the same residues in the alignment; “:” indicates conserved substitutions; “.” indicates semi-conserved substitutions; “-” indicates gaps in the alignment. (B and C) Immunoblotting in 293T cells expressing WT or mutant Twist1 treated with CHX (100 μg/ml) over a time course.

**Figure S4, related to Figure 6.**
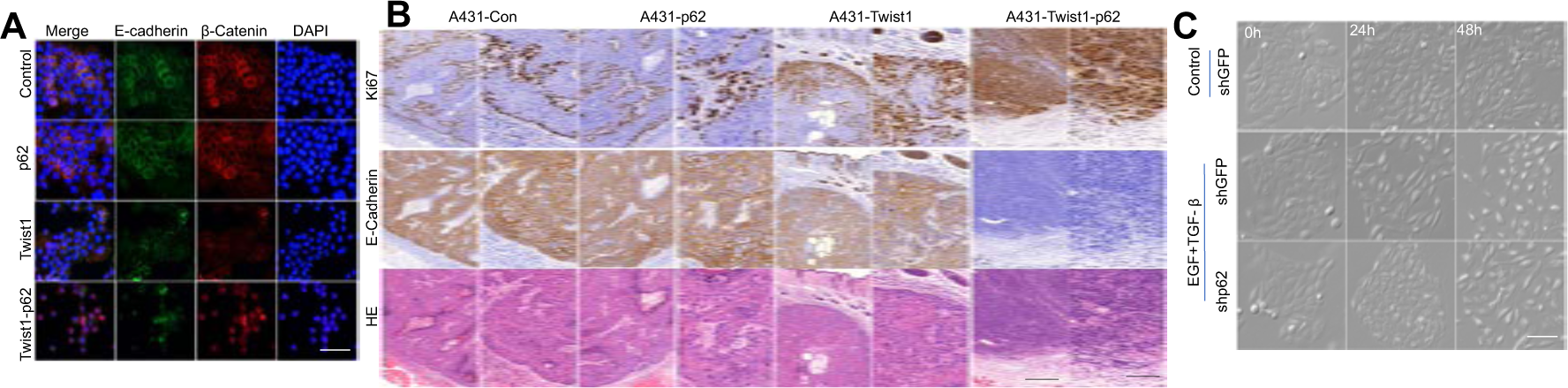
p62/Twist1 promotes tumor growth and metastasis. (A) Immunofluorescence staining of β-catenin in MEF cells with or without Atg5 deletion. Scale bar, 50 μm. (B) Histological and immunohistochemical staining in tumors. Scale bar, 200 μm and 50 μm for left and right panels, respectively. (C) Representative images of HaCaT cells with or without p62 knockdown, following treatment with or without EGF/TGF-β for 48 h. Scale bar, 50 μm.

